# The Impact of a Western Diet with High Salt on Metabolic Outcomes in Male C57bl/6J Mice

**DOI:** 10.1101/2025.10.17.683134

**Authors:** Michael E. Ponte, John C. Prom, Pradeep Devkota, Sireesha Yerrathota, Mea Carradine, Larry Ha, E. Matthew Morris, Andrew J. Lutkewitte

## Abstract

**Objective:** The Western diet promotes obesity and metabolic disease by increasing caloric intake and systemic inflammation. The typical Western diet is high in saturated fats, sugars, and salt. In pre-clinical rodent studies, the “Western” diet (also called the high-fat high-sucrose diet (HFHS)) is high in saturated fats and sugars (typically sucrose) but low in salt (<1% salt). As such, we sought investigate the impact of a chronic 3% NaCl Western diet (high-fat, high-sucrose + high salt (HFHS + Salt)) diet on systemic organ metabolism, liver mitochondrial function, and adipose tissue.

**Methods:** Thirty-six 8 week-old C57Bl/6J male mice were fed either a low-fat diet (LFD), a HFHS, or a HFHS + Salt diet for 16 weeks. Body weight, body composition, and food intake were monitored weekly. Glucose tolerance tests (GTT) and insulin concentrations were measured after 8 weeks of diet intervention to assess glucose and insulin homeostasis. Mice were euthanized at 16 weeks for liver mitochondrial respiration and tissue analysis.

**Results:** Over 16 weeks, the HFHS fed group gained significantly more weight than the other diet groups. Liver weights were similar in LFD and HFHS + Salt groups but higher in the HFHS group. Liver triglycerides (TAGs) were also similar between LFD and HFHS + Salt groups, while HFHS had elevated liver TAGs. Inguinal and brown adipose tissue depots were larger in both HFHS and HFHS + Salt vs. LFD. Surprisingly, the gonadal adipose tissue was significantly larger in the HFHS + Salt compared to HFHS and LFD groups – suggesting that a HFHS + Salt exacerbates gonadal adipose expansion more than typical rodent HFHS. Paradoxically, the addition of salt appears to have dampened expression of inflammation related genes (*Ccl2 & Adgre1*) in adipose tissue compared to HFHS alone. Metabolically, the HFHS+ Salt fed mice showed the highest glucose intolerance, followed by HFHS and then LFD groups. Liver mitochondrial respiration, assessed by changing ATP/ADP ratios, showed the HFHS group with the highest oxygen consumption, followed by HFHS + Salt, then LFD groups, highlighting differences in respiration with additional salt (HFHS vs HFHS + Salt).

**Conclusion:** While the excess salt mitigated some HFHS effects on weight gain and hepatic lipid accumulation, it exacerbated gonadal adipose expansion and impaired glucose tolerance. HFHS increased mitochondrial respiration, but salt addition appeared to dampen this effect. Dietary salt, within a high-fat/high-sucrose context, has differential impacts on metabolic outcomes compared to HFHS alone, underscoring the need for further research to fully understand how Western diets (high-fat, high-sucrose, *and high salt*) impact all aspects of metabolic health.

## 1. Introduction

The Western diet, typically rich in processed foods that contain added saturated fats, sugars (sucrose/fructose), and salt – promotes obesity and metabolic diseases not only by increasing caloric intake but also by inducing systemic inflammation and metabolic dysregulation [1, 2]. While the impact of these dietary components on systemic inflammation and metabolic dysregulation is recognized, the specific mechanisms, particularly concerning systemic and hepatic mitochondrial metabolism in response to dietary salt, remain poorly understood.

Dietary salt (sodium-chloride, NaCl) provides the body with sodium, essential for cellular homeostasis and various physiological functions, including maintaining extracellular fluid volume, osmolality, and maintaining membrane potentials through sodium/potassium exchange [3]. While only 1.25 g/day of salt is needed for normal physiological functions, modern diets significantly exceed this amount [4]. In the U.S., average daily sodium intake is over 3.2 g (equivalent to ∼8 g of table salt), and some populations consume up to ∼15-25 g/day [5, 6]. Excess salt intake is well-documented as a major contributor to cardiovascular diseases (CVD), one of the leading causes of death globally [7, 8]. Importantly, CVD is a major co-morbidity of obesity and metabolic syndrome, making the study of the complex/multi-component Western diet (high-fat, high-sugar, *and* high-salt) essential to understanding these interconnected public health crises. Global dietary guidelines recommending a maximum daily sodium intake of 2.3 g (5.75 g of table salt) [9]. The World Health Organization aims to reduce dietary salt (NaCl) intake to less than 5 g/day [10]. However, concerns have emerged regarding the potential downsides of strict sodium restriction and/or feasibility for individuals voluntarily reducing salt intake, sparking interest in salt replacement (potassium salt) therapy as potential alternatives [11, 12].

Recent studies suggest that salt may also influence metabolism and energy balance through mechanisms like increasing lipolysis, thermogenesis, and regulating key hormones such as leptin, natriuretic peptides, and aldosterone [13–16]. Both high and low salt intake have been associated with metabolic dysfunction, including insulin resistance, leptin resistance, obesity, and metabolic syndrome [13, 14, 17]. Conflicting findings on salt’s role in energy homeostasis may result from varying research designs in human and animal studies [17–21]. Clinic population studies such as the Trials of Hypertension Prevention (TOPH I & II) in the 1980s and subsequent follow ups in 2014/16 found a direct linear relationship between average daily sodium intake during the trial and cardiovascular disease risk. However, it’s unclear what the participants were eating in the years after the study – likely confounding the follow up results [22]. More recent studies have shown associations between high sodium and low potassium with increased cardiovascular risk – while increasing potassium intake may alleviate some risk [23]. Despite the well-established links between dietary salt and cardiovascular disease and emerging evidence of broader metabolic effects, the impact of excess dietary sodium on systemic metabolism and in particular hepatic mitochondrial function remains largely unexplored.

This gap in our understanding that various dietary amounts of dietary salt can have on metabolic outcomes highlights the need to better understand how excess dietary salt influences energy homeostasis and glucose metabolism, beyond its relatively well-known cardiovascular effects.

While excess sodium intake has been linked to insulin resistance and metabolic disease in human populations, high sodium in animal models has been shown to reduce glucose tolerance and insulin sensitivity, while low-sodium diets increase insulin-sensitizing adipokines and decrease adipose inflammation [5–9]. The effects of sodium on insulin sensitivity likely involve the renin-angiotensin system (RAS), which is linked to insulin resistance [24, 25]. Paradoxically, short-term sodium restriction has been shown to increase insulin resistance through RAS activation [20], while mineralocorticoid receptor blockade improves insulin sensitivity and reduces inflammation [26]. Sodium also influences immune cell behavior, promoting inflammatory responses that worsen metabolic outcomes [27–29].

Sodium’s role in fat deposition and obesity is relatively complex. High sodium intake is associated with obesity in humans [29–31], although animal studies show mixed results. Some research suggests sodium increases fat mass without affecting body weight [32], while other studies indicate it reduces both body weight and fat mass [33, 34]. In diet-induced obesity models, high sodium intake has been shown to prevent weight gain [35, 36].

The effects of chronic high-fat/high-sucrose/high-sodium (HFHS + Salt) diets on hepatic mitochondrial dynamics in the context of obesity are not well understood. Further, previous studies did not monitor on the week-to-week changes in food intake (as high-salt may change palatability) which could drive the differences in body composition

Therefore, this study aims to address this gap by investigating the direct impact of chronic Western diet consumption on hepatic mitochondrial function and metabolic outcomes. Better understanding these mechanisms will be crucial for developing more translatable preclinical dietary models to ultimately mitigate the metabolic consequences of excessive salt intake in the context of modern Western diets (high-fat, high-sucrose, and *high-salt*).

## 2. Materials & Methods

### Animals & Diet Paradigm

Animal protocols were approved by the Institutional Animal Care and Use Committee at the University of Kansas Medical Center. All experiments were carried out in accordance with the Guide for the Care and Use of Laboratory Animals. 36 male C57Bl/6J mice (#000664, Jackson Laboratory, Bar Harbor, ME, USA) were purchased from Jackson Laboratory at 6 weeks of age. Mice were acclimated to housing at ∼28⁰C on 12/12 reverse light cycle (dark 10:00 – 20:00), with ad libitum access to water and standard chow for 2 weeks prior to random group assignment [37]. Three diet groups were established: low-fat diet (10% kcal fat, 3.5% kcal sucrose, and 3.85 kcal/g energy density, Research Diets D12110704), high-fat/high-sucrose (45% kcal fat, 17% kcal sucrose, and 4.73 kcal/g energy density, Research Diets D12451), and high-fat/high-sucrose + 3% NaCl (Salt) (45% kcal fat, 17% kcal sucrose, and 4.73 kcal/g energy density Research Diets Special Diet # D06041001). On day 1 of the experiment, mice were given their respective diets for 16-weeks with fresh diet added weekly.

### Anthropometrics and energy intake

Body weight measurements started on Day 1 and were measured weekly. Food intake was assessed each week, and body composition was measured weekly using the EchoMRI-1100 system (EchoMRI, Houston, TX). Fat-free mass (FFM) was calculated as the difference between body weight (BW) and fat mass (FM). Week 10 body composition was not assessed due to equipment failure. Energy intake was calculated as the energy density of LFD (3.85 kcal/g), HFHS (4.73 kcal/g), or HFHS + Salt (4.6 kcal/g) times the food intake for 7 days.

### Glucose tolerance tests and insulin secretion assays

At week 8 of the diet intervention, animals had food withdrawn for 4-hours (at 8am) prior to a baseline blood glucose reading after which they were given an intraperitoneal (IP) injection of glucose (1.5 g/kg lean mass) to assess glucose homeostasis. Blood glucose and insulin were assessed from the tail vein at 0, 15, 30, 60, 90, and 120 minutes after IP injection. ∼50ul of serum was collected during each time point and frozen at −80°C. Insulin levels were assessed using a Mouse Insulin ELISA kit (Crystal Chem, Elk Grove Village, Illinois).

### Liver TAG Content and NAFL Activity Scoring

Liver TAGs and NAFL activity scoring were determined as previously described [38]. Briefly, ∼100 mg of frozen liver tissue was homogenized using a bead homogenizer in ice cold PBS (1mL/mg tissue). 100 µL of liver lysate was added to 100 µL of 1 % sodium deoxycholate the tubes were vortexed and heated at 37°C for 5 min to solubilize lipids. TAG content was measured enzymatically (Thermo Fisher TR22421). For NAFL scoring liver tissue was fixed in 10 % neutral buffered formalin for 48 hours and stored in 70 % ethanol until embedded and processed by The Kansas Intellectual and Developmental Disabilities Research Center (KIDDRC) Histology Core. H&E-stained liver sections were scored by an independent-blinded clinical pathologist using the Brunt Method [39].

### Mitochondrial Respiration

Liver mitochondria were isolated as previously described [40]. Briefly, ∼ 1 g of liver was homogenized (glass-on-teflon) in 8 mL of ice-cold mitochondrial isolation buffer (220 mM mannitol, 70 mM sucrose, 10 mM Tris, 1 mM EDTA, pH adjusted to 7.4 with KOH). The homogenate was centrifuged (4°C, 10 min, 1500 g), the supernatant was transferred to a round bottom tube, and centrifuged (4°C, 10 min, 8000x g). The pellet was resuspended in 6 mL of ice-cold mitochondrial isolation buffer using a glass-on-glass homogenizer, and centrifuged again (4°C, 10 min, 6000 x g). This final pellet was resuspended in ∼0.75 mL of modified mitochondrial respiration buffer (MiRO5) (0.5 mM EGTA, 3 mM MgCl2, 60 mM KMES, 20 mM glucose, 10mM KH2PO4, 20 mM HEPES, 110 mM sucrose, 0.1% BSA, 0.0625 mM free CoA, and 2.5mM carnitine, pH∼7.4). The protein concentration for both suspensions was determined by BCA assay.

### Liver Mitochondrial Creatine Kinase Clamp

Change in respiratory rate of isolated liver mitochondria during changes in Δ_G_ATP were performed as previously described with minor changes [40, 41]. Briefly, 2 mM malate, isolated liver mitochondria (50ul), 10 mM palmitoyl CoA, 10 mM palmitoyl-carnitine, 5 mM ATP, 5 mM creatine, and 20 U/mL creatine kinase were added to Oroboros chambers containing mitochondrial respiration buffer. ADP dependent respiration was initiated by the addition of 1 mM PCr, and sequential additions of PCr to 3, 6, 9, 12, 15, 18, 21, 24, 27, 30 mM reduced respiration toward baseline. The free energy of ATP for each PCr concentration was calculated as described [40]. The linear respiration data from 9 mM to 21 mM PCr was used and the conductance represented as the slope of the line. The data is oriented to represent a simulated increase in energy demand produced by the clamp.

### Liver Mitochondrial Palmitoyl-CoA Sensitivity

Briefly, oxygen consumption was measured during changes in palmitoyl-CoA concentrations (from 5uM to 30uM) after additions of 2mM malate, 2500U/mL hexokinase, and 500mM ADP.

### Adipose Tissue Diameter

Adipose tissue was fixed in 10 % neutral buffered formalin for 48 hours and stored in 70 % ethanol until embedded and processed by the KIDDRC Histology Core. H&E-stained adipose slides were scanned by the KUMC Biospecimen Repository Core Facility using a ZEISS Digital Slide Scanner Axioscan 7. Adipocyte diameter was measured using Zeiss Zen software by averaging 15 adipocytes of 3 separate slide sections per mouse.

### Gene Expression

Liver and kidney RNA was isolated from ∼ 25 mg of liver tissue using a RNeasy Plus Mini Kit (Qiagen, Valencia, CA, USA). The cDNA for liver tissues was produced using the ImProm-II RT system (Promega, Madison, WI, USA). Liver RT-PCR was performed using a Bio-Rad CFX Connect Real-Time System (Bio-Rad, Hercules, CA, USA) and SYBR Green. Adipose tissue RNA was isolated from ∼50 mg of tissue by homogenization in 1 mL of Qiazol (Qiagen). Adipose RNA was extracted using a RNeasy Lipid kit (Qiagen). Adipose cDNA was produced using a High-Capacity cDNA Reverse Transcription Kit (ThermoFisher). Gene specific values for liver were calculated using the using the −2^ΔΔct^ method and normalized to relative *Cyclophilin B* (*Ppib*, liver and kidney) or 36*b4* (adipose) mRNA expression values. Primer sequences are listed in **Table 1**.

**Table 1.**
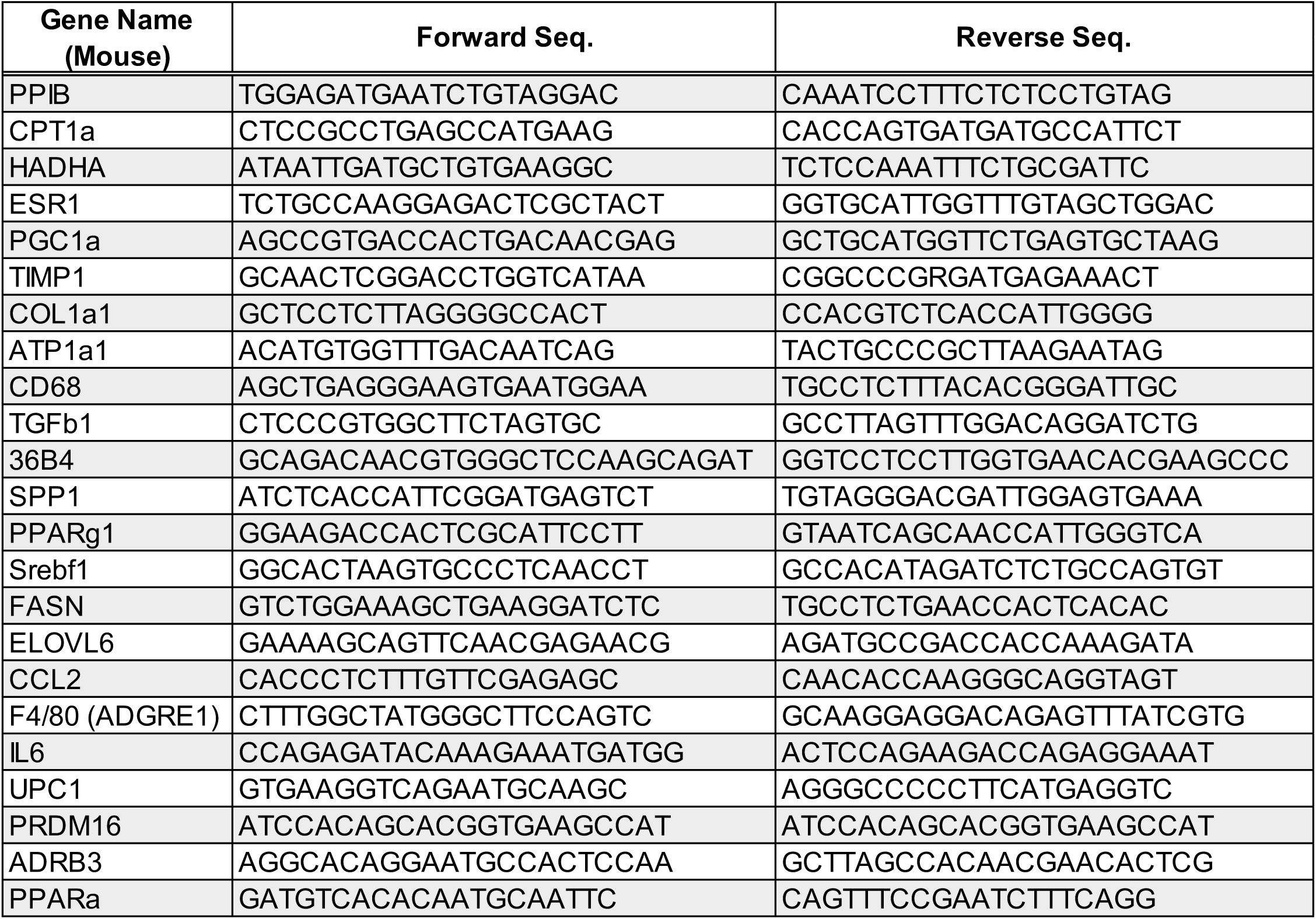
RT PCR primer sequences.

### Plasma metabolites

Plasma samples were analyzed using a Multiplex adipokine assay (Millipore-Sigma, MADKMAG-71K) by the CTBB core) for concentrations of IL-6, Leptin, MCP-1, PAI-1, Resistin, TNFα. Plasma triglycerides (TAGs), HDL, total cholesterol, AST and ALT were measured by IDEXX BioAnalytics (North Graphton, MA).

### Statistical Analysis

GraphPad Prism version 10.0 (GraphPad Software, San Diego, CA) was used for statistical analysis for each experiment unless otherwise mentioned. Data are presented as means and standard error. The two-standard deviation test was utilized to test for outliers within group. Data was assessed using one-way ANOVA with appropriate post-hoc tests for multi-group comparisons, or Student’s t-test for pairwise comparisons where indicated. Significance was considered as *p* > 0.05; LFD vs HFHS (γ), LFD vs HFHS + Salt (δ), HFHS vs HFHS + Salt (β).

## 3. Results

### 3.1. Dietary Salt Mitigates Overall Weight Gain but Worsens Glucose Tolerance

At the start of the study, all groups had similar body weights (∼22g). Over the 16-week intervention, the HFHS group gained significantly more body weight (∼22g) compared to the HFHS + Salt group (∼19g) and the LFD group (∼9g) (**Figure 1A**). This weight gain was primarily driven by a dramatic increase in fat mass, with only modest gains in lean mass (**Figure 1B & C**). Although weekly energy intake was initially elevated in HFHS and HFHS + Salt groups, around week 6 all the groups began to consume similar weekly kcals (**Figure 1D**). The cumulative energy intake was highest in the HFHS group, followed closely by the HFHS + Salt group, and lowest in the LFD group, generally aligning with the observed weight changes (**Figure 1D-F**). Notably, at dietary week 11, weekly energy intake was negatively affected by an institutional-mandated cage change. This caused a subtle loss in body weight of the HFHS and HFHS + Salt fed mice but not the LFD fed controls (**Figure 1** see arrows).

**Figure 1.**
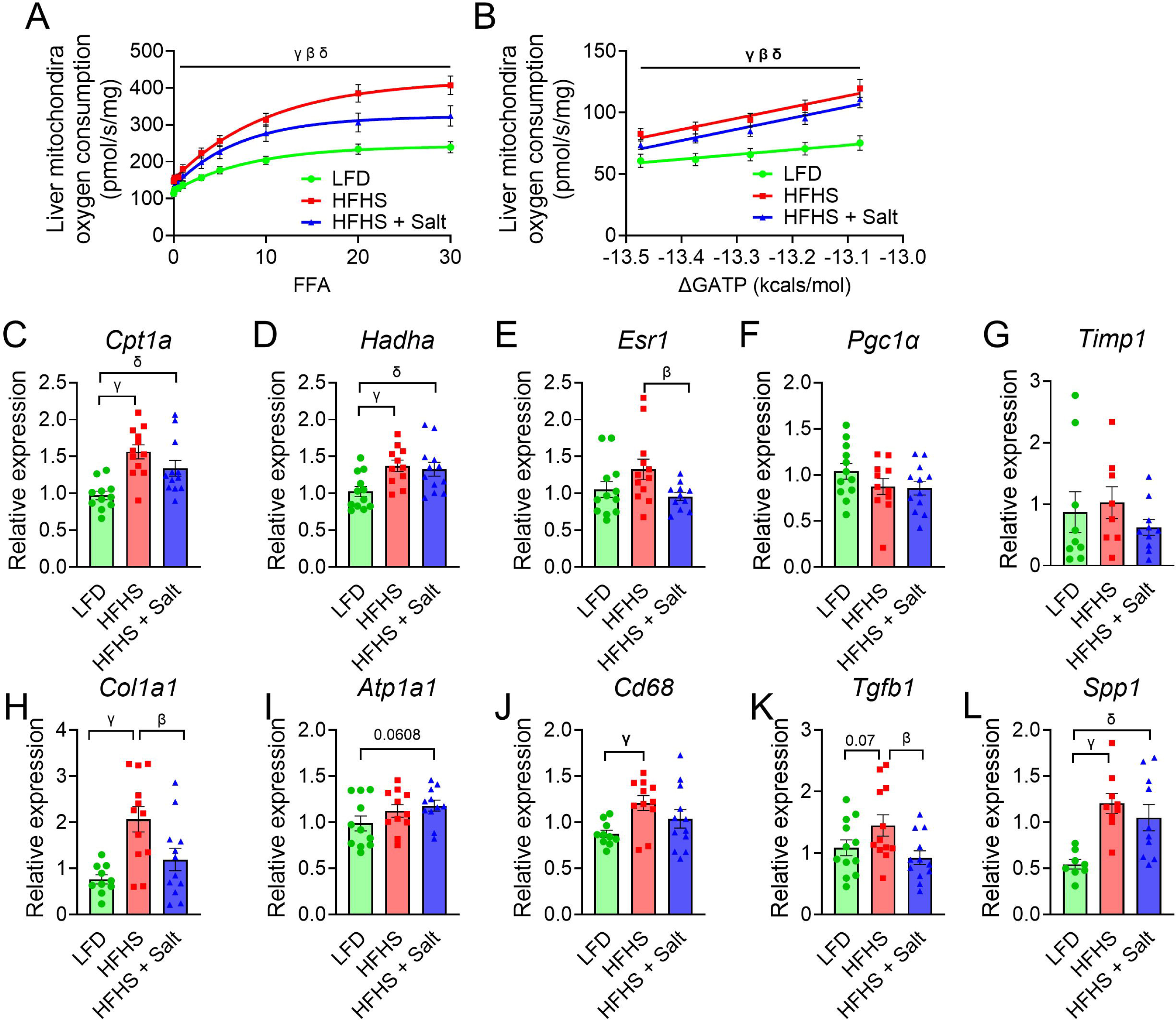
Metabolic and Body Composition Analysis after 16 Weeks of Diet Interventions. **A)** Body weight changes over 16 weeks during chronic diet interventions. **B)** Lean mass changes over 16 weeks during chronic diet interventions. **C)** Fat mass changes over 16 weeks during chronic diet interventions. **D)** Weekly energy intake in Kcal during the chronic diet interventions. **E)** Weekly energy intake normalized to body weight (Kcal/g) during the chronic diet interventions. **F)** Cumulative energy intake in Kcal during chronic diet interventions. In panels A-F, arrows indicate a “cage change” event. LFD: Low-fat diet; HFHS: High-fat, high-sugar diet; HFHS + Salt: High-fat, high-sugar diet with added salt. **Data presented as mean ± SEM. Statistical analysis was performed using one-way ANOVA with Fisher’s LSD post-hoc test.** Significance: *p* > 0.05; : LFD vs HFHS (γ), LFD vs HFHS + Salt (δ), HFHS vs HFHS + Salt (β).

Despite having lower overall body weight, mice on the HFHS + Salt diet exhibited the most severe glucose intolerance. During a glucose tolerance test (GTT) (1.5mg/kg glucose IP injection) at 8 weeks, the HFHS + Salt group displayed the highest blood glucose excursion, which was significantly greater than the LFD group as measured by the adjusted area under the curve (AAUC) (**Figure 2A & B**). This impaired glucose homeostasis was associated with a blunted insulin secretion response during the GTT in both HFHS and HFHS + Salt groups compared to the LFD group (**Figure 2C & D**).

**Figure 2.**
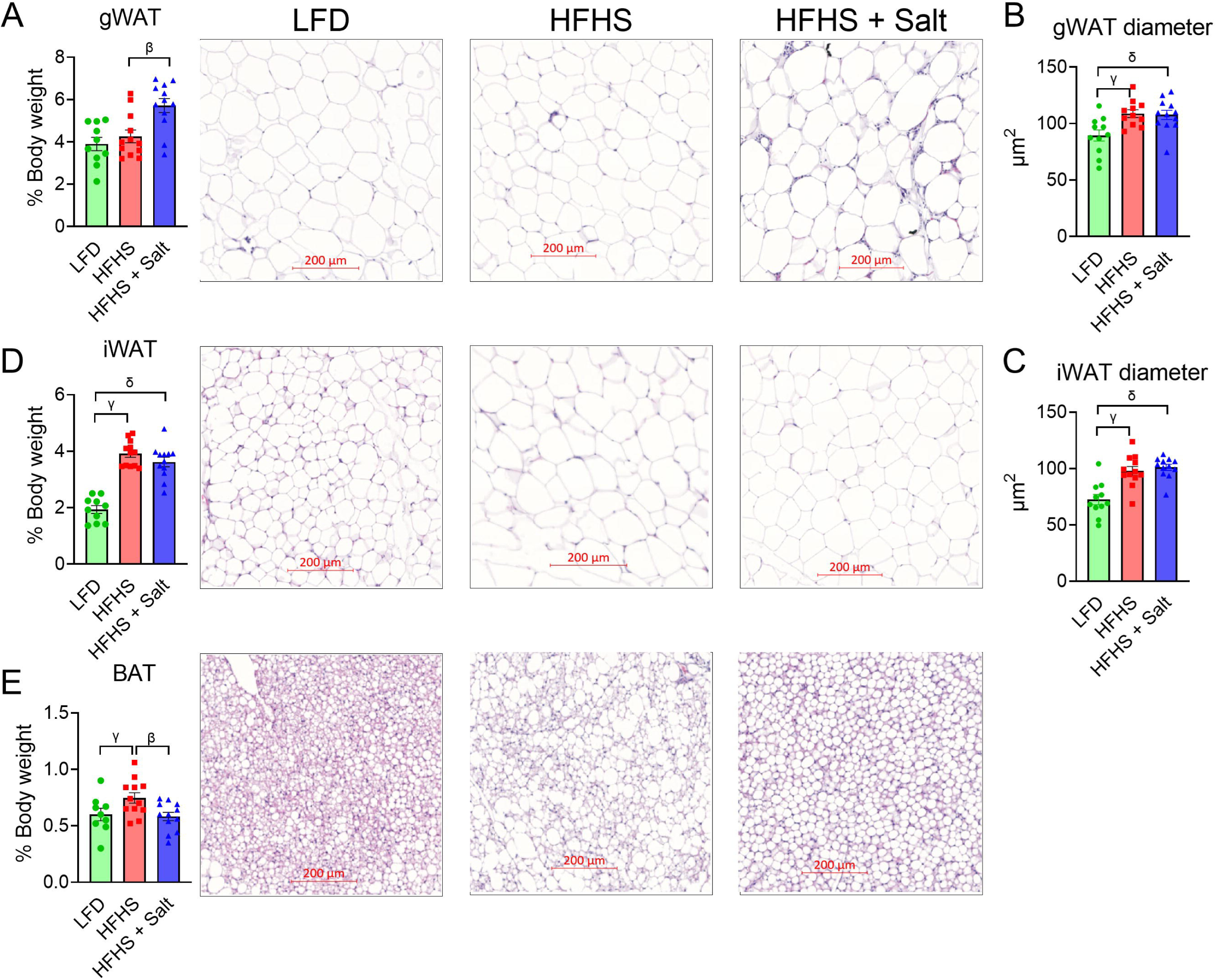
Glucose and Insulin Homeostasis. **A)** Blood glucose levels during a glucose tolerance test (GTT) following an intraperitoneal (IP) injection of 1.5 mg/kg glucose after 8 weeks of diet intervention. **B)** Adjusted area under the curve (AAUC) for the GTT. **C)** Insulin secretion during the GTT, **D)** expressed as % of initial value. LFD: Low-fat diet; HFHS: High-fat, high-sugar diet; HFHS + Salt: High-fat, high-sugar diet with added salt. Data presented as mean ± SEM. Statistical analysis was performed using one-way ANOVA with Fisher’s LSD post-hoc test. Significance *p* > 0.05; LFD vs HFHS (γ), LFD vs HFHS + Salt (δ), HFHS vs HFHS + Salt (β).

### 3.2. Excess Dietary Salt May Protect Against Hepatic Steatosis on a High-Fat Diet

To determine the extent of metabolic derangement caused by HFHS and HFHS + Salt feeding, we analyzed a major metabolic organ the liver. The HFHS diet group saw elevated markers of hepatic steatosis, as evidenced by a significant increase in liver weight and an increase in liver triglycerides (TAGs) (**Figure 3A & B**). This was corroborated by significantly elevated NAFL activity scores (NAS), driven primarily by steatosis (**Figure 3D-G**). Interestingly, spleen weight (as a % body weight), a surrogate for portal hypertension, was similarly decreased in both high-fat high-sucrose groups relative to LFD – suggesting no/minimal portal vein hypertension occurred in these groups (**Figure 3H**). HFHS feeding caused a significant increase in serum ALT but not AST, a marker of liver damage (**Figure 3I, J**). Remarkably, the addition of 3% salt to the HFHS diet dampened these effects; the HFHS + Salt group had liver weights, TAG content, and AST/ALT levels that were more like the LFD group (**Figure 3A, B, D-G, I & J**). While additional dietary salt may have “protected” the liver from excessive lipid accumulation, both high-fat diets induced systemic dyslipidemia. Serum HDL and total cholesterol were significantly elevated in both HFHS and HFHS + Salt groups compared to LFD (**Figure 3L & M**).

**Figure 3.**
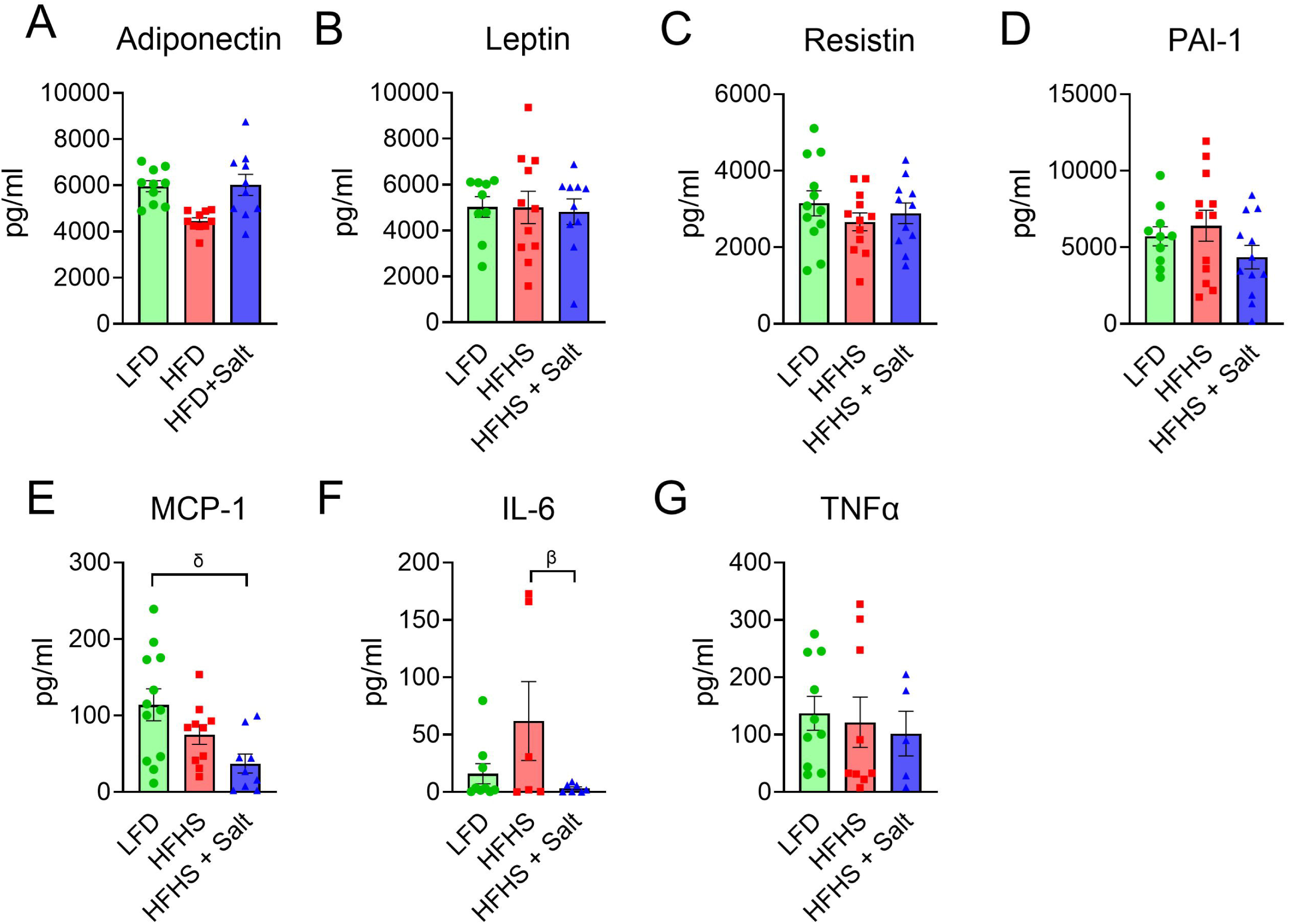
Hepatic and Systemic Metabolic Markers after 16 Weeks of Diet Interventions. **A)** Liver weight expressed as a percentage of total body weight. **B)** Liver triglyceride (TAGs) content (mg/g liver tissue) after 16 weeks of diet intervention. **C)** Representative (10x) Hematoxylin and Eosin (H&E) stained images of liver tissue from each diet group. **D)** Composite NAS (NAFLD Activity Score) score, derived from summing scores for **E)** ballooning degeneration. **F)** lobular inflammation. **G)** steatosis. **H)** Spleen weight expressed as a percentage of total body weight. **I)** Serum Aspartate Aminotransferase (AST) levels (U/L). **J)** Serum Alanine Aminotransferase (ALT) levels (U/L). **K)** Serum triglyceride (TAGs) levels (mg/dL). **L)** Serum High-Density Lipoprotein (HDL) levels (mg/dL). **M)** Total serum cholesterol levels (mg/dL). LFD: Low-fat diet; HFHS: High-fat, high-sugar diet; HFHS + Salt: High-fat, high-sugar diet with added salt. Data presented as mean ± SEM. Statistical analysis was performed using one-way ANOVA with Fisher’s LSD post-hoc test. Significance *p* > 0.05; LFD vs HFHS (γ), LFD vs HFHS + Salt (δ), HFHS vs HFHS + Salt (β).

### 3.3. HFHS Diet Increases Liver Mitochondrial Respiration, an Effect Dampened by Dietary Salt

To further investigate the mechanisms behind the hepatic phenotype, we assessed liver mitochondrial function. Hepatic mitochondria isolated from the HFHS group exhibited the highest rate of oxygen consumption in response to increasing concentrations of the fatty acid palmitoyl-CoA, indicating a heightened capacity for fatty acid oxidation (**Figure 4A**). A similar pattern was observed using a creatine kinase clamp to assess respiration across a range of ATP free energy (ΔGATP) values – going from low-to-high (state of rest to exercise) energy demand (**Figure 4B**). In contrast, the HFHS + Salt group showed an intermediate respiratory phenotype, with oxygen consumption rates significantly lower than the HFHS group but higher than the LFD group (**Figure 4A & B**).

**Figure 4.**
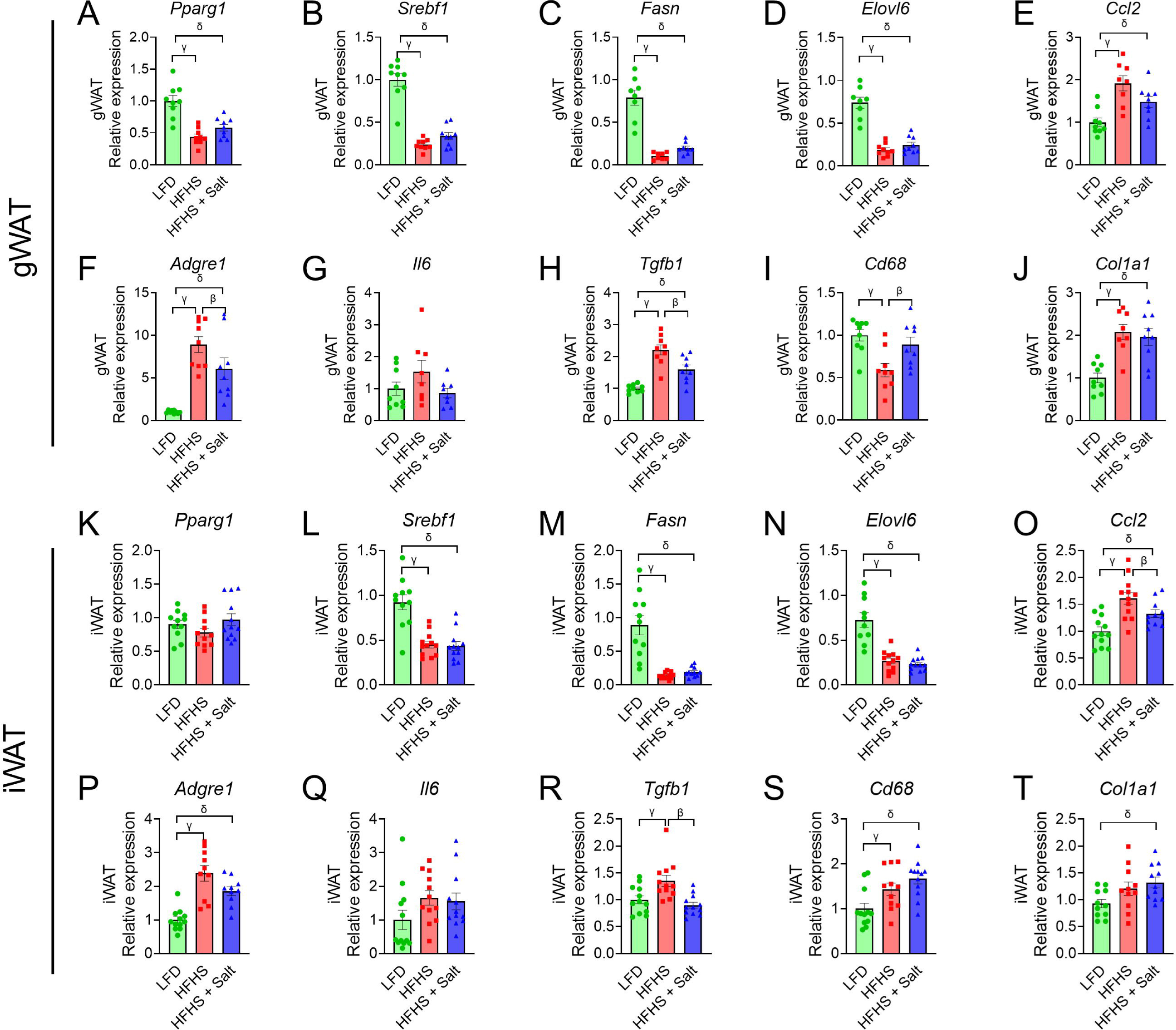
Liver Mitochondrial Respiration and Gene Expression. **A)** Oxygen consumption (pmol/s/mg protein) of isolated liver mitochondria in response to increasing concentrations of Palmitoyl-CoA. **B)** Oxygen consumption (pmol/s/mg protein) of isolated liver mitochondria across a range of ATP free energy (ΔG ATP) values, maintained via a creatine kinase clamp. **C-L)** Relative mRNA expression levels in liver tissue for genes involved in metabolism & inflammation: **C)** *Carnitine palmitoyltransferase 1a* (*Cpt1a*), **D)** *Hydroxyacyl-CoA dehydrogenase alpha* (*Hadha*), **E)** *Estrogen receptor 1* (*Esr1*), **F)** *Peroxisome proliferator-activated receptor gamma coactivator 1-alpha* (*Pgc1α*), **G)** *Tissue inhibitor of metalloproteinases 1* (*Timp1*), **H)** *Collagen type I alpha 1 chain* (*Col1a1*), **I)** *ATPase Na+/K+ transporting subunit alpha 1* (*Atp1a1*), **J)** *Cluster of differentiation 68* (*Cd68*), **K)** *Transforming growth factor beta 1* (*Tgfb1*), and **L)** *Secreted phosphoprotein 1* (*Spp1*). LFD: Low-fat diet; HFHS: High-fat, high-sugar diet; HFHS + Salt: High-fat, high-sugar diet with added salt. Data presented as mean ± SEM. Statistical analysis was performed using one-way ANOVA with Fisher’s LSD post-hoc test**, non-linear fit, and simple linear regression**. Significance *p* > 0.05; LFD vs HFHS (γ), LFD vs HFHS + Salt (δ), HFHS vs HFHS + Salt (β).

Gene expression analysis of the liver revealed that markers of fatty acid oxidation (*Cpt1a*, *Hadha*) were upregulated in both high-fat groups compared to LFD (**Figure 4C & D**). However, genes associated with fibrosis (*Col1a1*), inflammation (*Cd68*), and tissue remodeling (*Spp1*, *Tgfb1*) were most highly expressed in the HFHS group, and their expression was reduced by the addition of dietary salt (**Figure 4H-L**). Interestingly, expression of the sodium-potassium pump subunit *Atp1a1* showed a trend towards being highest in the HFHS + Salt group (p=0.068) (**Figure 4I**).

### 3.4. Excess Dietary Salt Alters Adipose Tissue Distribution

In stark contrast to its effects on liver and overall body weight, excess dietary salt specifically exacerbated the expansion of gonadal (visceral) fat. The gonadal white adipose tissue (gWAT) depot was significantly larger in the HFHS + Salt group compared to both the HFHS and LFD groups (**Figure 5A**). This was accompanied by a significant increase in gWAT adipocyte diameter compared to the LFD group, although gWAT diameter was similar between the HFHS + Salt and HFHS groups (**Figure 5B**). Inguinal (subcutaneous) white adipose tissue (iWAT) depots and adipocyte size were similarly increased in both high-fat groups compared to LFD (**Figure 5C & D**). In contrast, brown adipose tissue (BAT) was largest in the HFHS group, an effect that was normalized by the additional 3% dietary salt (**Figure 5E**).

**Figure 5.**
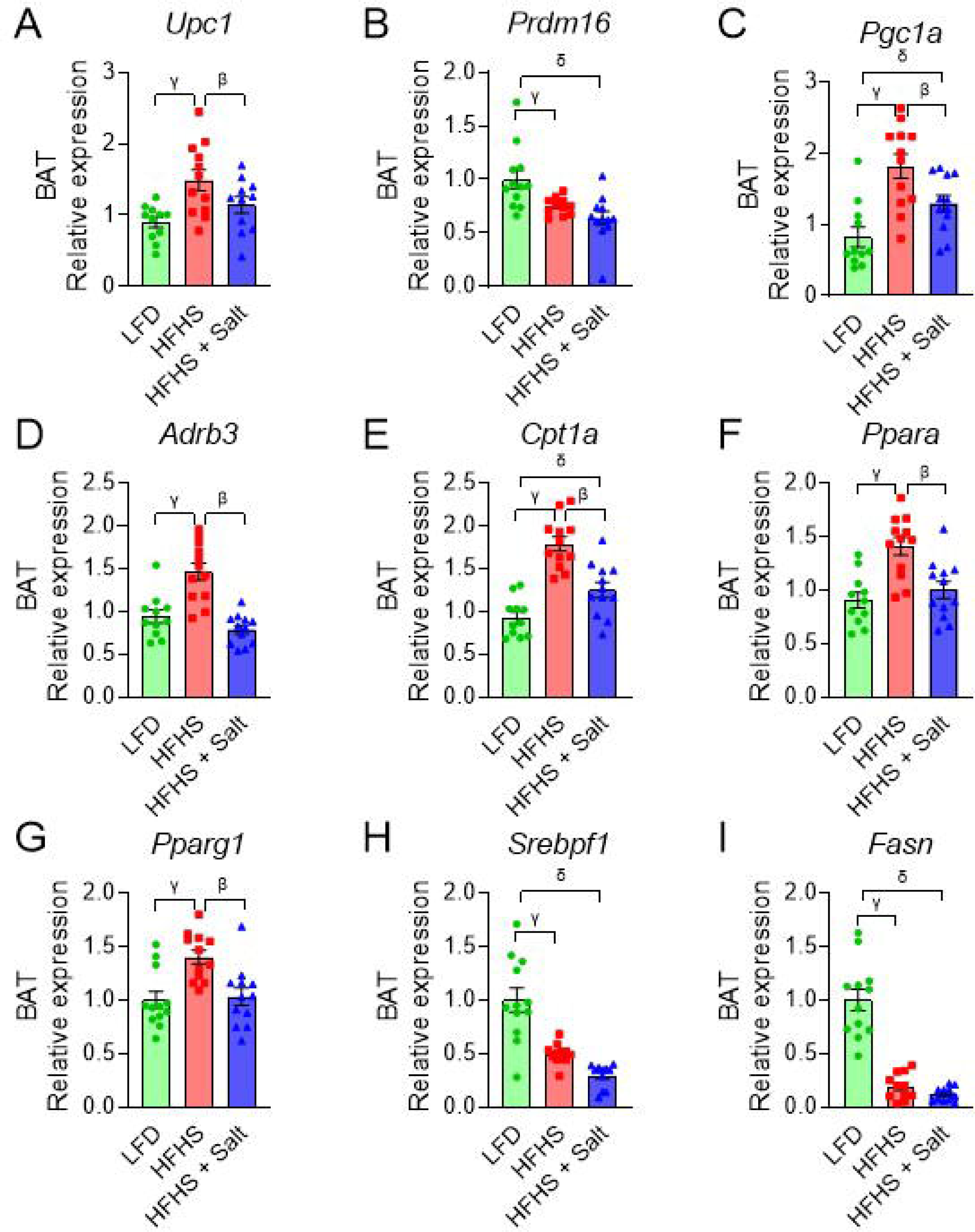
Adipose Tissue Depot Analysis. **A)** Gonadal white adipose tissue (gWAT) weight expressed as a percentage of total body weight, with representative Hematoxylin and Eosin (H&E) stained images for each diet group. **B)** gWAT diameter. C) Inguinal white adipose tissue (iWAT) weight expressed as a percentage of total body weight, with representative H&E-stained images for each diet group. **C)** iWAT diameters. **E)** Brown adipose tissue (BAT) weight expressed as a percentage of total body weight, with representative H&E stained images for each diet group. LFD: Low-fat diet; HFHS: High-fat, high-sugar diet; HFHS + Salt: High-fat, high-sugar diet with added salt. Data presented as mean ± SEM. Statistical analysis was performed using one-way ANOVA with Fisher’s LSD post-hoc test. Significance *p* > 0.05; LFD vs HFHS (γ), LFD vs HFHS + Salt (δ), HFHS vs HFHS + Salt (β).

### 3.5. HFHS Diets Induce Adipose Inflammation, with Salt Addition Exerting Modulatory Effects

Plasma adipokine levels of adiponectin, leptin, resistin, and plasminogen activator inhibitor 1 (PAI-1) were not different among the groups (**Figure 6A-D**). Plasma MCP-1 was significantly lower in the HFHS + Salt group compared to LFD group (**Figure 6E**). The pro-inflammatory cytokine IL-6 was significantly elevated in the HFHS group, an effect that was completely abolished in the HFHS + Salt group while TNF-α was unchanged (**Figure 6F, G**).

**Figure 6.**
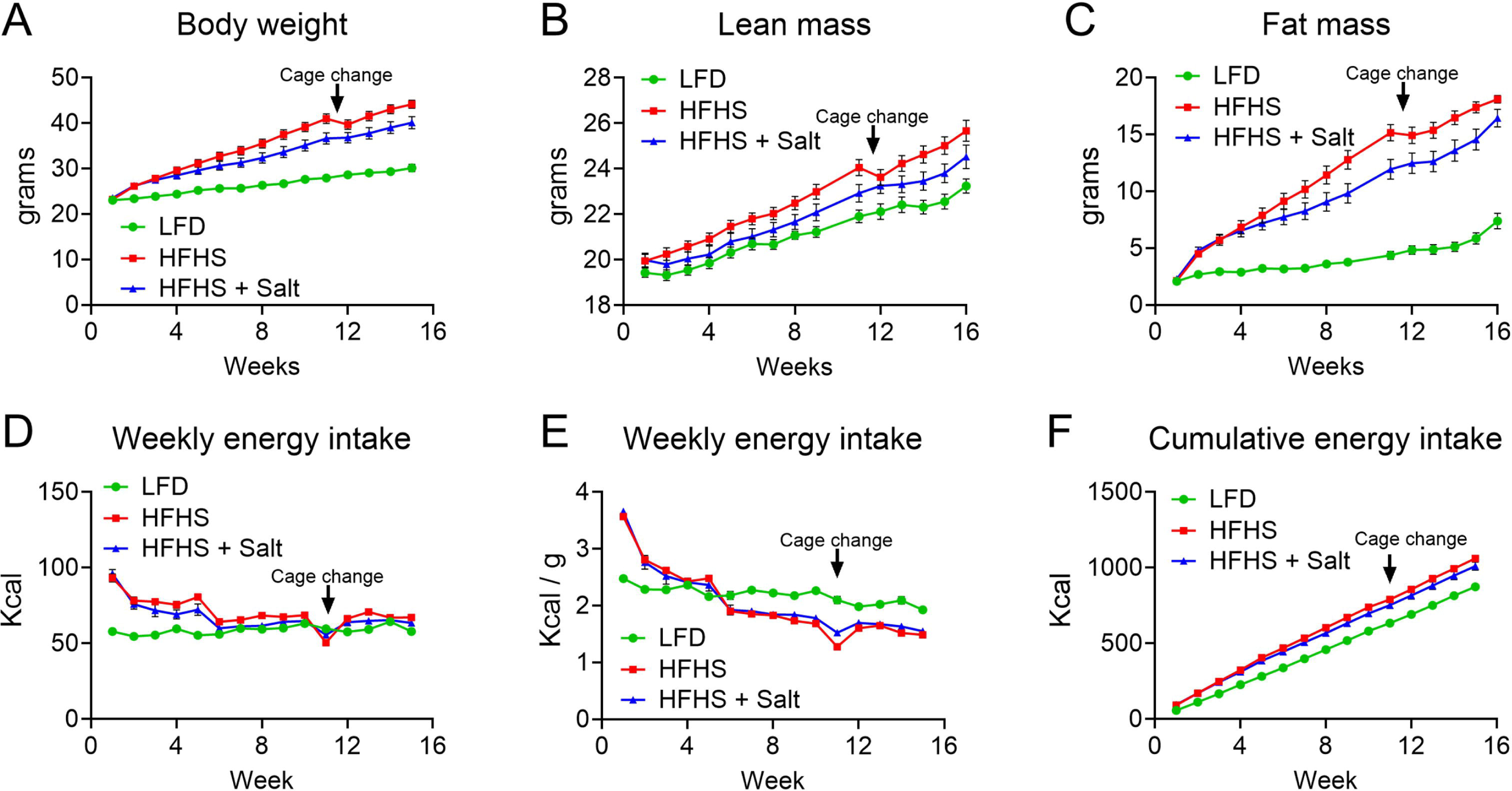
Circulating Adipokine and Inflammatory Cytokine Levels. Plasma concentrations (pg/ml) of: **A)** Adiponectin, **B)** Leptin, **C)** Resistin, **D)** Plasminogen Activator Inhibitor-1 (PAI-1), **E)** Monocyte Chemoattractant Protein-1 (MCP-1), **F)** Interleukin-6 (IL-6), and **G)** Tumor Necrosis Factor alpha (TNFα). LFD: Low-fat diet; HFHS: High-fat, high-sugar diet; HFHS + Salt: High-fat, high-sugar diet with added salt. Data presented as mean ± SEM. Statistical analysis was performed using one-way ANOVA with Fisher’s LSD post-hoc test. Significance *p* > 0.05; LFD vs HFHS (γ), LFD vs HFHS + Salt (δ), HFHS vs HFHS + Salt (β).

In gWAT, several genes involved in adipocyte function (fat synthesis and storage), *Pparg1*, *Srebf1*, *Fasn*, and *Elovl6*, are all significantly upregulated in the LFD group compared to the HFHS and HFHS + Salt groups (**Figure 7A-D**). In contrast to the lipogenic genes, the inflammatory and fibrotic markers *Ccl2* (a monocyte chemoattractant), *Adgre1* (a macrophage marker), and *Tgfb1* (a gene involved in inflammation and fibrosis) are all significantly increased in the HFHS group compared to LFD – while the expression in the HFHS + Salt group is intermediary (**Figure 7 E-H**). Interestingly, *Cd68* (macrophage marker) was significantly elevated in the LFD and HFHS + Salt compared to the HFHS group (**Figure 7I**). *Col1a1* (fibrosis marker) was significantly lower in the LFD group compared to the HFHS and HFHS + Salt groups (**Figure7J**).

**Figure 7.**
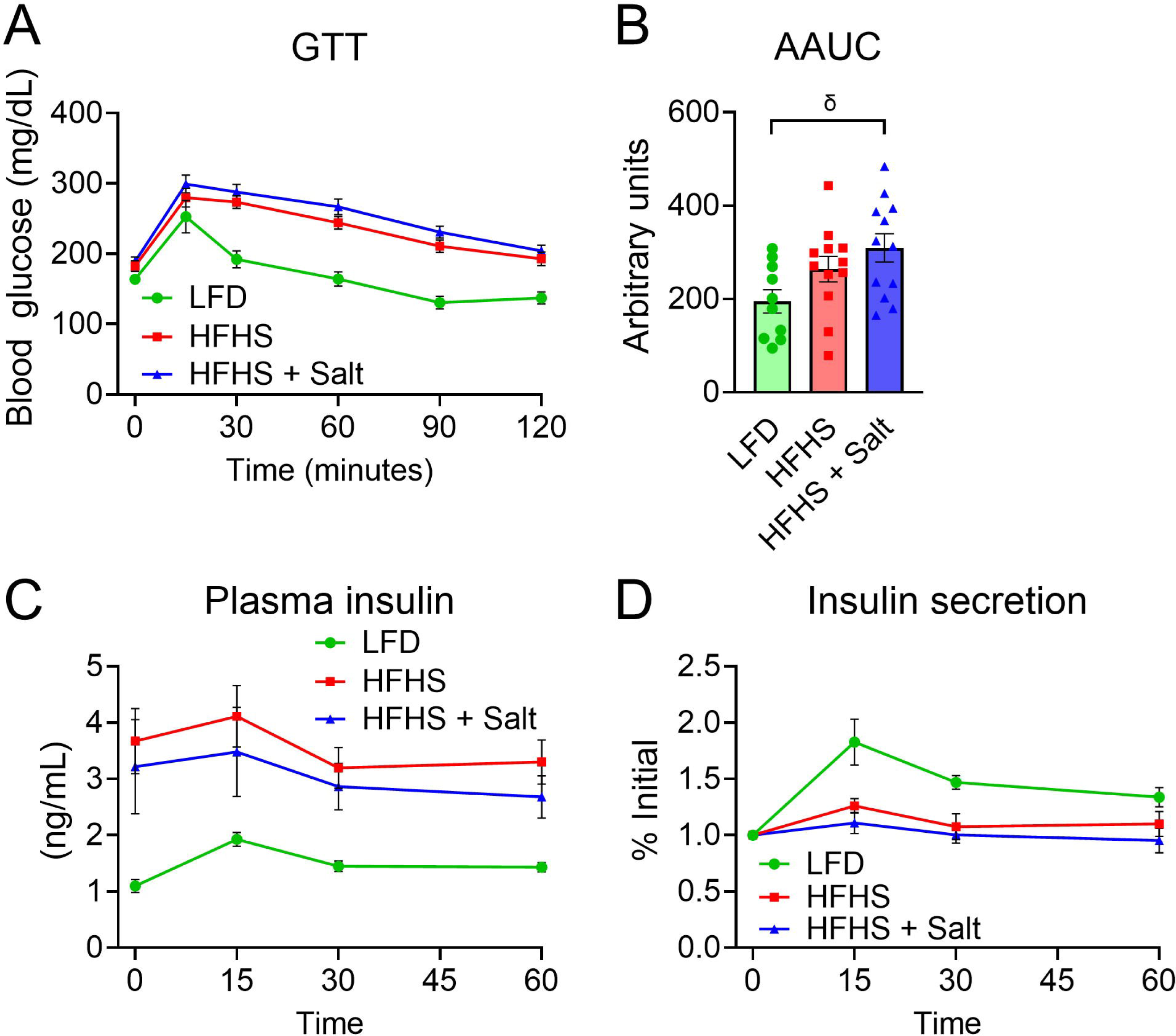
Adipose Tissue Gene Expression. Relative mRNA expression levels in gonadal white adipose tissue (gWAT) for genes involved in lipogenesis, inflammation, and fibrosis including: **A)** *Peroxisome proliferator-activated receptor gamma 1* (*Pparg1*), **B)** *Sterol regulatory element-binding protein 1* (*Srebf1*), **C)** *Fatty acid synthase* (*Fasn*), **D)** *Elongation of very long chain fatty acids protein 6* (*Elovl6*), **E)** *C-C motif chemokine ligand 2* (*Ccl2*), **F)** *Adhesion G protein-coupled receptor E1* (*Adgre1*), **G)** *Interleukin 6* (*Il6*), **H)** *Transforming growth factor beta 1* (*Tgfb1*), **I)** *Cluster of Differentiation 68* (*Cd68*), and **J)** *Collagen type I alpha 1 chain* (*Col1a1*). Relative mRNA expression levels in inguinal white adipose tissue (iWAT) for: **K)** *Pparg1*, **L)** *Srebf1*, **M)** *Fasn*, **N)** *Elovl6*, **O)** *Ccl2*, **P)** *Adgre1*, **Q)** *Il6*, **R)** *Tgfb1*, **S)** *Cd68*, and **T)** *Col1a1*. LFD: Low-fat diet; HFHS: High-fat, high-sugar diet; HFHS + Salt: High-fat, high-sugar diet with added salt. Data presented as mean ± SEM. Statistical analysis was performed using one-way ANOVA with Fisher’s LSD post-hoc test. Significance *p* > 0.05; : LFD vs HFHS (γ), LFD vs HFHS + Salt (δ), HFHS vs HFHS + Salt (β).

iWAT gene expression revealed no significant differences in *Pparg1* expression between groups, however, *Srebf1*, *Fasn*, and *Elovl6* showed significant increases in relative expression of LFD animals compared the HFHS groups (**Figure 7K-N**). Both *Ccl2* and *Adgre1* were significantly elevated in the HFHS and HFHS + Salt groups – with the Salt mice having and intermediary expression between the HFHS and LFD groups (**Figure 7O & P**). Like the gWAT *Il6* expression – there were no significant differences between the groups, although the HFHS and HFHS + Salt trended higher in iWAT tissue (**Figure 7Q**). Also, like gWAT, iWAT expression of *Tgfb1* was significantly elevated in the HFHS group compared to LFD and HFHS + Salt (**Figure 7R**). Interestingly, *Cd68* expression was elevated in both HFHS and HFHS + Salt groups compared to LFD, and likewise for *Col1a1* expression (**Figure 7S & T**).

Brown adipose tissue gene analysis revealed increased expression of *Upc1* (thermogenesis) in the HFHS group compared to LFD and HFHS + Salt groups (**Figure 8A**). *Prdm16* (BAT induction and differentiation) was significantly elevated in the LFD group compared to both HFHS groups (**Figure 8B**). Genes involving in transcriptional regulation of energy balance and metabolism (*Pgc1a*, *Adrb3*, *Cpt1a*, *Ppara*, and *Pparg1*) showed significantly increased expression in the HFHS group compared to LFD and HFHS + Salt groups (**Figure 8C-G**). While *Srebpf1* and *Fasn*, also lipid metabolism genes, showed significantly increased in expression in the LFD group compared with both HFHS groups (**Figure 8H & I**). Together these data highlight the depot specific changes in response to our diet interventions.

**Figure 8.**
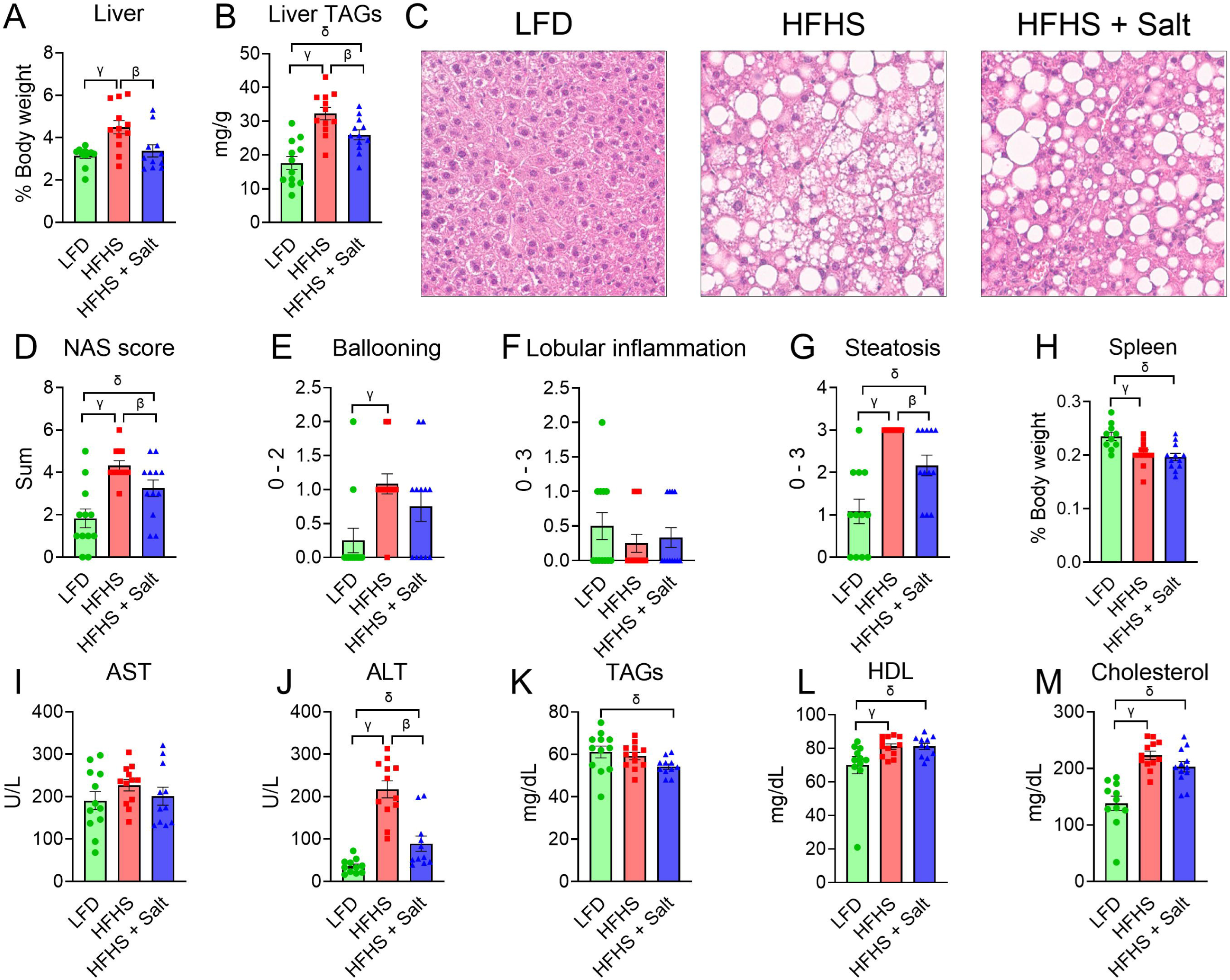
Brown Adipose Tissue Gene Expression. Relative mRNA expression levels in brown adipose tissue (BAT) for genes involved in thermogenesis, fatty acid oxidation, and lipogenesis including: **A)** *Uncoupling protein 1* (*Ucp1*), **B)** *PR/SET domain 16* (*Prdm16*), **C)** *Peroxisome proliferator-activated receptor gamma coactivator 1-alpha* (*Pgc1a*), **D)** *Adrenoceptor beta 3* (*Adrb3*), **E)** *Carnitine palmitoyltransferase 1a* (*Cpt1a*), **F)** *Peroxisome proliferator-activated receptor alpha* (*Ppara*), **G)** *Peroxisome proliferator-activated receptor gamma 1* (*Pparg1*), **H)** *Sterol regulatory element-binding protein 1* (*Srebf1*), and **I)** *Fatty acid synthase* (*Fasn*). LFD: Low-fat diet; HFHS: High-fat, high-sugar diet; HFHS + Salt: High-fat, high-sugar diet with added salt. Data presented as mean ± SEM. Statistical analysis was performed using one-way ANOVA with Fisher’s LSD post-hoc test. Significance *p* > 0.05; LFD vs HFHS (γ), LFD vs HFHS + Salt (δ), HFHS vs HFHS + Salt (β).

## 4.1 Discussion

This study aimed to investigate the impact of a chronic high-fat/high-sucrose diet with and without added salt (to mimic a *modern* Western diet) on metabolic outcomes in male C57B/6J mice. Our findings demonstrate differential effects on body composition, glucose metabolism, liver function, hepatic mitochondrial respiration, and adipose tissue distribution. With increasing healthcare burdens related to chronic Western diets (high fat, sugar, *and* salt) and associated metabolic dysfunction, it is critical we better understand how these dietary compositions impact metabolic outcomes compared to traditional high-fat, high-sugar (primarily sucrose and often low salt) diets commonly used in preclinical research models [42–44].

Our results demonstrate that while both HFHS and HFHS + Salt diets led to increased body weight compared to the low-fat diet (LFD), the HFHS group accumulated more weight over the 16-week period compared to the HFHS + Salt group (**Figure 1A**). This difference in weight gain was primarily driven by increased fat mass accumulation, with the HFHS group exhibiting greater total fat mass (**Figure 1C**). Surprisingly, despite HFHS + Salt having statistically similar cumulative energy intake to the HFHS group (**Figure 1F**), the added salt appears to have mitigated some of the weight-promoting effects of the high-fat/high-sucrose diet. We acknowledge that the cumulative energy intake is slightly lower in the Salt fed group compared to the HFHS diet alone. Since even a slight decrease in energy intake can reduce weight gain over an extended period, this could explain the reduction in weight gain and therefore many of the metabolic observations here. Further research to determine the optimal % of NaCl to avoid palatability aversion yet still drive hypertension and metabolic disease.

Although the additional salt mostly improved the metabolic outcomes measured in this study, there was a striking change in adipose tissue distribution (**Figure 5**) – the HFHS + Salt diet led to an increase in gonadal adipose tissue, which is consistent with previous work [45]. Surprisingly, the addition of salt did not exacerbate adipose inflammation. In fact, the salt lowered tissue and plasma markers of inflammation in gWAT and iWAT (**Figures 5 and 6**). Inflammatory and fibrotic markers were consistently upregulated in the high-fat diet groups. In gWAT, *Ccl2*, *Adgre1*, and *Tgfb*1 were all significantly increased in the HFHS group compared to LFD, with the HFHS + Salt group showing an intermediary expression (**Figure 7E-H**). Similarly, in iWAT, both *Ccl2* and *Adgre1* were significantly elevated in the HFHS and HFHS + Salt groups, confirming a broad inflammatory response in both groups (**Figure 7O & P**). Interestingly, *Cd68* expression was lower in the HFHS group compared to both LFD and HFHS + Salt groups in gWAT but elevated in iWAT unlike the LFD group (**Figure 7I & S**). This was accompanied by a significant increase in *Col1a1* expression in both HFHS and HFHS + Salt groups compared to LFD in gWAT (**Figure 7J**). While both high-fat diets induced fibrosis (*Col1a1*), the addition of salt “dampened” the expression of *Tgfb1* in both iWAT and gWAT (**Figure 7H & R**). Surprisingly, plasma IL-6 levels were significantly elevated in the HFHS group but completely normalized in the HFHS + Salt group – this suggests that the added salt might have a modulating effect on the inflammatory response, potentially by affecting the production or clearance of IL-6 (**Figure 6F**). Plasma MCP-1 levels were significantly lower in the HFHS + Salt group compared to LFD, suggesting excess dietary salt influences chemokine signaling (**Figure 6E**). In contrast, brown adipose tissue (BAT) weight was significantly increased in the HFHS group compared to HFHS + Salt and LFD (**Figure 5E**). The mechanisms underlying these specific, depot-dependent effects of high dietary salt on adipose tissue warrant further investigation into adipocyte proliferation/expansion during high-salt exposure.

The addition of 3 % NaCl did not change spleen weight (a marker of portal hypertension, **Figure 3H**) nor heart or kidney weight (data not shown), suggesting that 3% NaCl + HFHS diet may not be sufficient to drive portal hypertension in C57Bl/6J male mice. Many hypertension studies use diets with as much as ∼4-8% NaCl; however, these studies often do not rigorously assess metabolic functions [46–48].

Increased liver triglycerides and elevated alanine aminotransferase (ALT) levels, hallmarks of hepatic steatosis and liver damage, were observed in the HFHS group (**Figure 3A, B & J**). However, the addition of salt to the HFHS diet appeared to ameliorate these effects, with liver triglyceride and ALT levels in the HFHS + Salt group being similar to the LFD group (**Figure 3A, B & J**). This suggests a potential “*protective*” or “*blunting*” effect of high-salt against hepatic lipid accumulation and liver injury markers in the context of a chronic high-fat/high-sucrose diet. This observation is consistent with the significantly lower liver weight observed in the HFHS + Salt group compared to the HFHS group, and its similarity to LFD (**Figure 3A**). Detailed scoring for markers of liver injury, including the overall NAFL activity score (NAS) and its components (ballooning, lobular inflammation, and steatosis), further revealed elevated scores, particularly for steatosis, in the HFHS group compared to LFD and HFHS + Salt (**Figure 3D-G**). Serum HDL and total cholesterol were significantly elevated in both high-fat, high-sucrose groups compared to LFD, suggesting that the added salt didn’t “*blunt*” circulating markers of dyslipidemia compared to HFHS alone (**Figure 3L & M**). One possibility for the improvement in liver metabolism is that lipids are repartitioned from the liver to the expanded gonadal adipose depot, where lipids are physiologically stored.

Metabolically, the HFHS + Salt group exhibited the highest degree of glucose intolerance, followed closely by the HFHS group, with the LFD group demonstrating the best glucose tolerance (**Figure 2A & B**). The absolute area under the curve (AAUC) for the GTT was significantly increased in the HFHS + Salt group compared to the LFD but not the HFHS group (**Figure 2B**). Fasting insulin was higher in both the HFHS and HFHS + Salt diet groups and insulin secretion as a response to the glucose bolus was blunted both high-fat-high sucrose groups. (**Figure 2C, D**). Future studies should include insulin tolerance testing and more detailed assessment of insulin kinetics to better characterize the effects of a true Western diet on insulin sensitivity and secretion dynamics in chronic high-fat fed mice.

Mitochondrial respiration studies revealed that the HFHS group exhibited the highest oxygen consumption rates in response to both increasing Palmitoyl-CoA (PCoA) concentrations and changes in ATP free energy via the creatine kinase (CK) clamp protocols (**Figure 4A & B**). While the HFHS + Salt group also showed elevated oxygen consumption compared to the LFD group, its magnitude was intermediate between the LFD and HFHS groups for both protocols. These findings suggest that both high-fat/high-sucrose diets, with or without salt, increase liver mitochondrial respiration, potentially reflecting increased fatty acid oxidation capacity or overall metabolic demand. Liver mRNA expression analysis revealed that markers of fatty acid oxidation (Cpt1a, Hadha) were upregulated in both high-fat groups, consistent with increased fatty acid metabolism capacity (**Figure 4C & D**). We also observed a trend towards increased Atp1a1 expression in the HFHS + Salt group (p=0.068), suggesting a potential mechanism where the increased sodium load from the diet elevates the energy demand of the Na+/K+-ATPase, thereby increasing energy expenditure and reducing the substrate available for storage in the liver.

## 4.2 Summary

In conclusion, this study demonstrates that the addition of dietary salt (NaCl) to a high-fat/high-sucrose diet has complex and sometimes paradoxical effects on metabolic parameters. While salt appears to mitigate some of the negative effects of a high-fat/high-sucrose diet on overall weight gain, hepatic lipid accumulation, liver injury markers (e.g., ALT), and serum IL-6 levels, it exacerbates gWAT expansion and impairs glucose tolerance. These findings highlight the importance of considering the combined and nuanced effects of individual dietary components within the context of a modern Western diet (high fat, high sugar (sucrose), *and* high salt). Our data emphasizes the need for further research to fully interpret the metabolic effects of a true Western diet in preclinical models. Future studies will focus on exploring the specific mechanisms by which salt influences adipose tissue metabolism, hepatic lipid accumulation, and glucose/insulin homeostasis.

